# HMMER Cut-off Threshold Tool (HMMERCTTER): Supervised Classification of Superfamily Protein Sequences with a reliable Cut-off Threshold

**DOI:** 10.1101/130443

**Authors:** Inti Anabela Pagnuco, María Victoria Revuelta, Hernán Gabriel Bondino, Marcel Brun, Arjen ten Have

## Abstract

Protein superfamilies can be divided into subfamilies of proteins with different functional characteristics. Their sequences can be classified hierarchically, which is part of sequence function assignation. Typically, there are no clear subfamily hallmarks that would allow pattern-based function assignation by which this task is mostly achieved based on the similarity principle. This is hampered by the lack of a score cut-off that is both sensitive and specific.

HMMER Cut-off Threshold Tool (HMMERCTTER) adds a reliable cut-off threshold to the popular HMMER. Using a high quality superfamily phylogeny, it clusters a set of training sequences such that the cluster-specific HMMER profiles show 100% precision and recall (P&R), thereby generating a specific threshold as inclusion cut-off. Profiles and threshold are then used as classifiers to screen a target dataset. Iterative inclusion of novel sequences to groups and the corresponding HMMER profiles results in high sensitivity while specificity is maintained by imposing 100% P&R. In three presented case studies of protein superfamilies, classification of large datasets with 100% P&R was achieved with over 95% coverage. Limits and caveats are presented and explained.

HMMERCTTER is a promising protein superfamily sequence classifier provided high quality training datasets are used. It provides a decision support system that aids in the difficult task of sequence function assignation in the twilight zone of sequence similarity. A package containing source code and full dataset will be deposited at Github and is available for reviewers at: https://www.dropbox.com/s/aacao6ggcak30bg/Repo.tar.gz?dl=0

**Author summary:** The enormous amount of genome sequences made available in the last decade provide new challenges for scientists. An important step in genome sequence processing is function assignation of the encoded protein sequences, typically based on the similarity principle: The more similar sequences are, the more likely they encode the same function. However, evolution generated many protein superfamilies that consist of various subfamilies with different functional characteristics, such as substrate specificity, optimal activity conditions or the catalyzed reaction. The classification of superfamily sequences to their respective subfamilies can be performed based on similarity but since the different subfamilies also remain similar, it requires a reliable similarity score cut-off.

We present a tool that clusters training sequences and describes them in profiles that identify cluster members with higher similarity scores than non-cluster members, i.e. with 100% precision and recall. This defines a score cut-off threshold. Profiles and thresholds are then used to classify other sequences. Classified sequences are included in the profiles in order to improve sensitivity while maintaining specificity by imposing 100% precision and recall. Results on three case studies show that the tool can correctly classify complex superfamilies with over 95% coverage.

HMMERCTTER is meant as a decision support system for the expert biologist rather than the computational biologist.

## Introduction

Protein sequence function annotation is one of the major tasks of computational genomics. The most widely applied tools are based on the similarity principle: the higher the similarity between sequences, the higher the probability these have the same function. For instance BLAST [1]□ is a search machine, routinely used by biologists in order to identify the function of their query sequences. Although the functional protein sequence space conforms only a minor part of the polypeptide sequence space, similarity based sequence annotation is hampered by many problems.

The requirement of a set of reliably annotated sequences is fundamental and despite the steady development of UniProt [2]□, incorrectly annotated sequences form a major obstacle in sequence function annotation. A second problem is that of the high sequence variation that apparently is allowed for large numbers of protein families. The resulting sensitivity problem becomes more apparent when protein superfamilies are considered. The evolution of proteins has for a large part been instigated by gene duplications and the resulting process of functional redundancy and diversification [3]□. Paralogs can obtain novel functions due to relaxed functional constraints, often while maintaining its original function. All together this results in intricate superfamilies where function annotation by similarity scoring is hampered by problems of sensitivity and specificity combined with imperfect annotation of reference sequences. This problem increases when taking into account the fact that, in the post genome era, biologists want to obtain annotations at the subfamily level, rather than the superfamily level. In other words, queries are performed to identify orthologs rather than homologs. Pattern based search strategies, such as provided by, for instance, Prosite [4]□ can be applied to increase specificity, as for instance when combined with BLAST [5]□.

HMMER [6]□ is another tool for sequence function annotation. Rather than comparing a query with a reference sequence database, it uses a database of mathematical profiles that describe multiple sequence alignments of protein families. Besides that the higher information usage generates a higher sensitivity, profile databases are more easily curated and improved curation results in significantly improved annotation. Several web servers that use HMMER to search their profile databases exist, all based on different objectives and principles. Pfam [7]□ is a collection of domain profiles whereas Superfamily [8]□ describes proteins based on the Structural Classification of Proteins [9]□. Although both platforms show moderate levels of hierarchical organization, the objective of these major function annotation tools is to annotate at a superfamily rather than a subfamily level.

A more recent trend is that of phylogenomics [10]□ and hierarchical sub-clustering of superfamilies. Phylogenomics is based on the correct idea that phylogeny allows for a better clustering than similarity since it is based on evolutionary models. Since phylogeny is a form of hierarchical clustering, it also allows for the identification of subfamilies. A number of algorithms and phylogenomics-based sequence annotation platforms have been developed in the last two decades. RIO [11]□ is dedicated to the identification of orthologs and paralogs using bootstrapping to calculate a confidence value for orthology. SCI-PHY [12]□ and GEMMA [13]□ use agglomerative clustering. SCI-PHY implements a subfamily encoding cost minimization to define sub-clusters whereas GEMMA’s sub-clustering is based on E-value. The coding cost minimization algorithm from SCI-PHY has been applied to Pfam and the identified subfamilies were analyzed for function shifts by means of the identification of Specificity Determining Positions (SDPs), resulting in the FunShift database [14]□. GEMMA was used to subcluster the CATH-Gene3D resource resulting in FunFHMMER [15]□. Here an automated cut-off was provided by a functional coherence index, also based on SDPs. CDD [16]□ is a domain database that includes an automated hierarchical classification of subfamilies, based on Bayesian analysis of what basically constitute SDPs [17]□. Sifter [18,19]□ uses an empirical Bayesian approach to combine function evidence from for the Gene Ontology Annotation (GOA) [20] database with a phylogenetic tree. Panther [21]□ is a database with curated protein families that, besides GOA data and phylogenetic trees, includes manually curated metabolic pathways. Hence, basically phylogenomics platforms apply SDPs, either computationally predicted or empirically identified, to obtain more specific partitions.

Since the ultimate goal of the above mentioned methods is to provide HMMER databases that cover functional protein space, they depend on heuristics. Partitions and classifications might therefore contain errors when compared with the real phylogeny. Furthermore, most of the methods are fully automated, which can result in clusters that do not correspond with functional clusters as determined by experts, the identified clusters being either too small or too large. More importantly however, is the fact that current sequence function annotation methods lack a cut-off that results both in high precision and in high recall. Certain platforms, such as Pfam, use curated trusted thresholds providing high precision. Others, such as Panther and CDD, are redundant and show either the hit with best E value or all significant hits. Combining high specificity with high sensitivity is arguably problematic.

Interestingly, the high information usage of HMMER has principally been deployed to increase sensitivity whereas in principle high information content can also be applied to increase specificity. Basically, HMMER aligns a sequence to an MSA and computes a score of residue-profile correspondence. The variation among the sequences, which take part in the underlying MSA, affects the score of a query-profile alignment. Sites with high information content, i.e. highly conserved sites, will give either high rewards or high penalties whereas sites with low information content, i.e. highly variable sites, will hardly contribute to the total score. Thus, a HMMER profile made from a variable superfamily-MSA will be less specific than a HMMER profile made from a conserved subfamily-MSA. Thus, representing large, complex superfamilies by the various subfamilies’ HMMER profiles will result in higher specificity, while presumably maintaining high sensitivity. Based on this principle, we developed a semi-automated, user-supervised procedure and pipeline that splits a superfamily into component subfamilies with the primary objective to cluster and classify its sequences with high P&R. Using a high quality phylogeny, HMMER Cut-off Threshold Tool or HMMERCTTER automatically identifies monophyletic sequence clusters that have 100% precision and recall (P&R) in a HMMER screening. In other words, it generates profiles made from cluster-specific MSAs that by *hmmsearch* identify all the cluster’s sequences with a score higher than that of any other sequence provided by the training set. This cluster-specific score threshold provides a specific inclusion cut-off. Subsequently, these clusters can be accepted or rejected by the user assisted with information presented in *hmmsearch* score plots. In the classification phase, target sequences are classified using searches with the cluster-specific HMMER profiles and the established cut-off threshold as classifiers. Both profiles and corresponding cut-offs are iteratively updated upon the inclusion of novel sequences during an automated and a subsequent user-controlled classification, while imposing 100% P&R.

The pipeline, which connects various existing softwares, is briefly described and demonstrated by detailed case studies of the alpha-crystallin domain (ACD) protein, the polygalacturonase (PG) and the phospholipase C (PLC) superfamilies. In the near future, HMMERCTTER will be extended towards the analysis of complete proteomes.

## Results

### Design of Method and Pipeline

Fig 1 outlines the HMMERCTTER training procedure and target sequence analysis, which are described in brief below and in detail in S Appendix 1. The training sequences are clustered using a user provided phylogeny. All possible monophyletic clusters are determined, sorted by size and tested as follows. The cluster’s sequences are aligned and used to generate a HMMER profile that is subsequently used to screen the cluster’s sequences as well as all training sequences. Obtained HMMER scores are compared and 100% P&R is obtained when the lowest scoring cluster sequence has a higher score than the highest scoring non-cluster sequence. 100% P&R clusters are provisionally accepted whereas non 100% P&R clusters are automatically rejected, An interface showing score plots of cluster and training sequences as well as a tree with the provisional clustering is presented as shown in S Fig 1A, at which point the user can reject or accept the cluster. Upon rejection, the program proceeds with the next cluster on the size-ordered list. Upon acceptance of a cluster, all its nested and overlapping clusters are removed from the list, and the program proceeds with the next cluster in the sorted list until no more clusters are encountered. This yields a number of clusters that show 100% P&R in HMMER profiling as well as, possibly, a number of unclustered orphan sequences.

**Fig 1:**
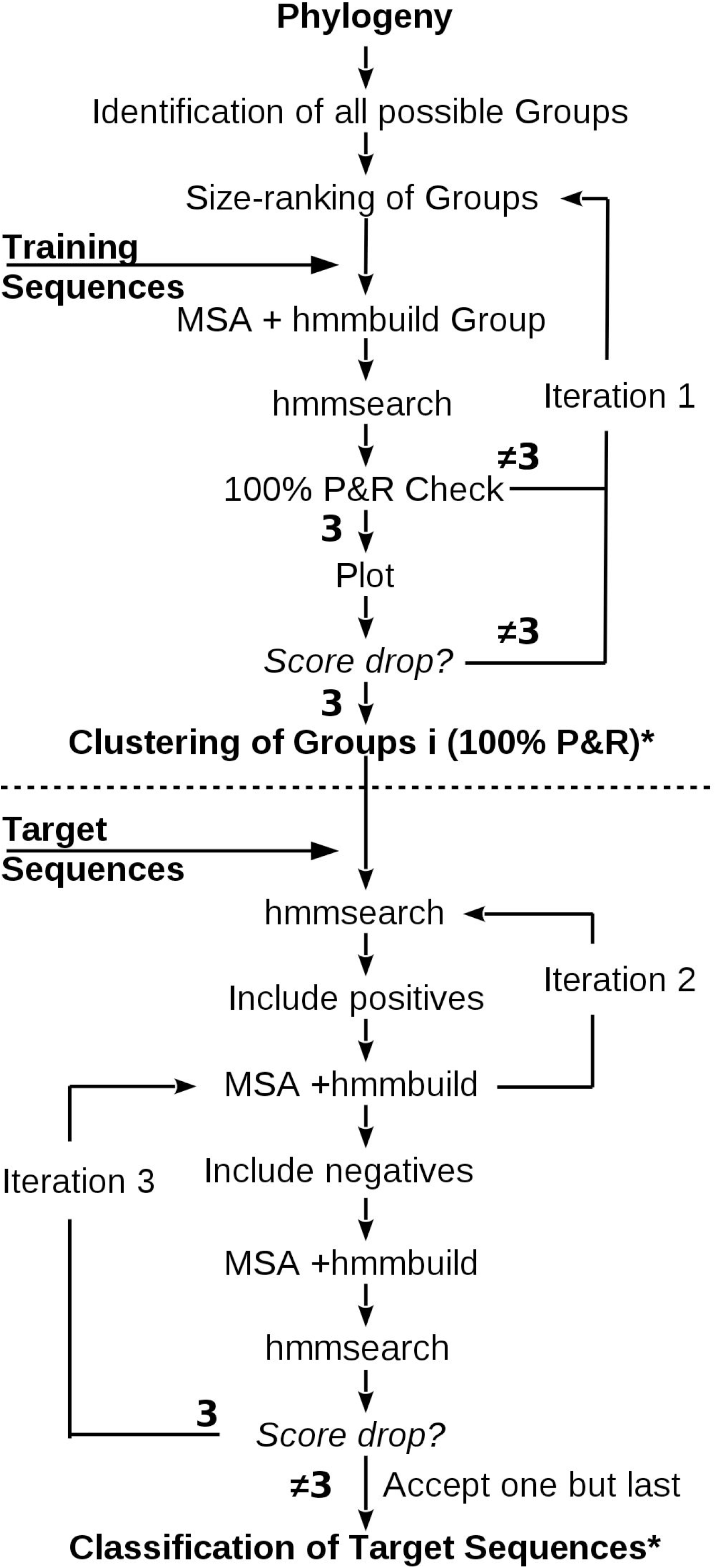
Flowchart of HMMERCTTER Pipeline. Training and target phase are separated by the dotted line. Monophyletic clusters of the training set are tested for 100% P&R, in descending size order. Iteration 1 is performed when a group is not accepted (either automatically or by user intervention) and the procedure is repeated with a smaller monophyletic group until no more groups are available for analysis. Accepted groups with corresponding HMMER profile and specific cut-off, defined by the 100% P&R rule, are used later to classify target sequences. Automated iteration cycle 2 is performed upon inclusion of sequences with prior 100% P&R. Upon convergence and user acceptance, supervised iteration 3 includes seemingly negatives upon a test for posterior 100% P&R, i.e. upon construction of a novel profile. Note that iteration 2 is nested inside iteration 3, albeit user controlled.* indicates that final clustering and classification do not necessarily show 100% coverage.

The HMMER profiles and corresponding cut-off scores, which equal the lowest scores of the clusters’ sequences, form the initial classifiers that are used for screening the target dataset. In order to clarify whether we refer to the clustering or the classification phase of the pipeline, a cluster results from the clustering phase whereas a group is the result of the classification phase. Sequences with scores equal or above the cluster threshold are automatically accepted and added to the cluster, forming a group. We refer to these sequences as prior positives since they were not yet included in the cluster or group when tested. Sequences are realigned to construct a new HMMER profile with a new cut-off score in order to obtain higher sensitivity in subsequent HMMER profiling. As such, groups remain 100% P&R provided classification overlap is prevented. When a target sequence becomes classified by more than one group, all groups are excluded from subsequent iterations. Conflicting training sequences are removed from all but the original group whereas conflicting target sequences are removed from all groups and target dataset.

This automated step of classification terminates upon data convergence, when no novel sequences with a score above the threshold are identified. Hitherto, all accepted sequences were accepted based on a prior inclusion HMMER cut-off threshold, i.e. by a HMMER profile that did not include the to be accepted sequence(s). However, certain sequences might only be accepted once their information has been included into the profile, i.e. according to a posterior inclusion HMMER cut-off threshold. Hence, in the subsequent classification step, sequences with a score below the threshold are considered. Candidates are included in the group and tested with a novel HMMER profile that includes the candidate. An interactive interface (S Fig 1B) allows the user to guide this process while 100% P&R remains imposed and classification conflicts remain prohibited as described for the automated phase. The process is terminated by the user, resulting in updated groups and a file that indicates which sequences generated conflicts.

### Algorithm Performance

We set out to test the pipeline using three protein superfamilies. The major objective was to identify putative problems and limits of HMMERCTTER and to survey the general applicability of the procedure. In all cases classification was performed with optimal coverage as primary criterion. Manual override during classification (i.e. rejecting group updates) was applied only when the drop in the HMMER score decreased by at least an order of magnitude. Performance was measured as coverage of corresponding sequence space. The first case consists of a published dataset that therefore restricts the sequence space to that of the published phylogeny, resulting in a fixed reference dataset with known classification. The other cases consist of novel datasets in which the sets of target sequences are formed by all homologous sequences identified from large reference proteomes dataset as described in materials and methods. In these cases no clear reference clustering exists and coverage is expressed as Sequence Space Coverage Interval which is the interval between the lowest and highest possible coverage of each group, restricted by monophyly.

### The plant ACD protein superfamily: A complex case with paraphyletic groups and repeats

Alpha crystallin domain (ACD) proteins form a large superfamily that include various subfamilies of the well described small heat shock proteins (sHSPs) as well as a number of poorly or not described subfamilies [22]□. Recently we identified 824 ACD proteins in 17 plant proteomes, using cluster-specific HMMER profiles manually made using a training set consisting of all ACD sequences identified in seven complete plant proteomes [22]□. This suggested the existence of 24 major and five minor subfamilies alongside two orphan sequences. Approximately half of the subfamilies are sHSPs, which functional classification is largely based on the cellular component of function. The remaining clusters include a family of transcriptional regulators, a family of salt stress induced proteins and 11 subfamilies of Uncharacterized ACD Proteins (UAP) [22]□. We took the datasets as previously used [22]□ but removed sequences of the five minor subfamilies and a single orphan, all distant sequences that fall inside the major sHSP-C1 cluster and prevent detection of the sHSP-C1 cluster since the algorithm imposes monophyly.

We obtained a sequence clustering that is nearly identical to the described functional classification (Fig 2A). The major discrepancy consists of UAPVII that was not 100% P&R and further divided into groups 11, 14, 19 and 24. In order to determine how the several groups behave in HMMERCTTER classification we compared its classification with the final reference phylogeny of the complete sequence set, under the assumption that the phylogeny is correct.

**Fig 2:**
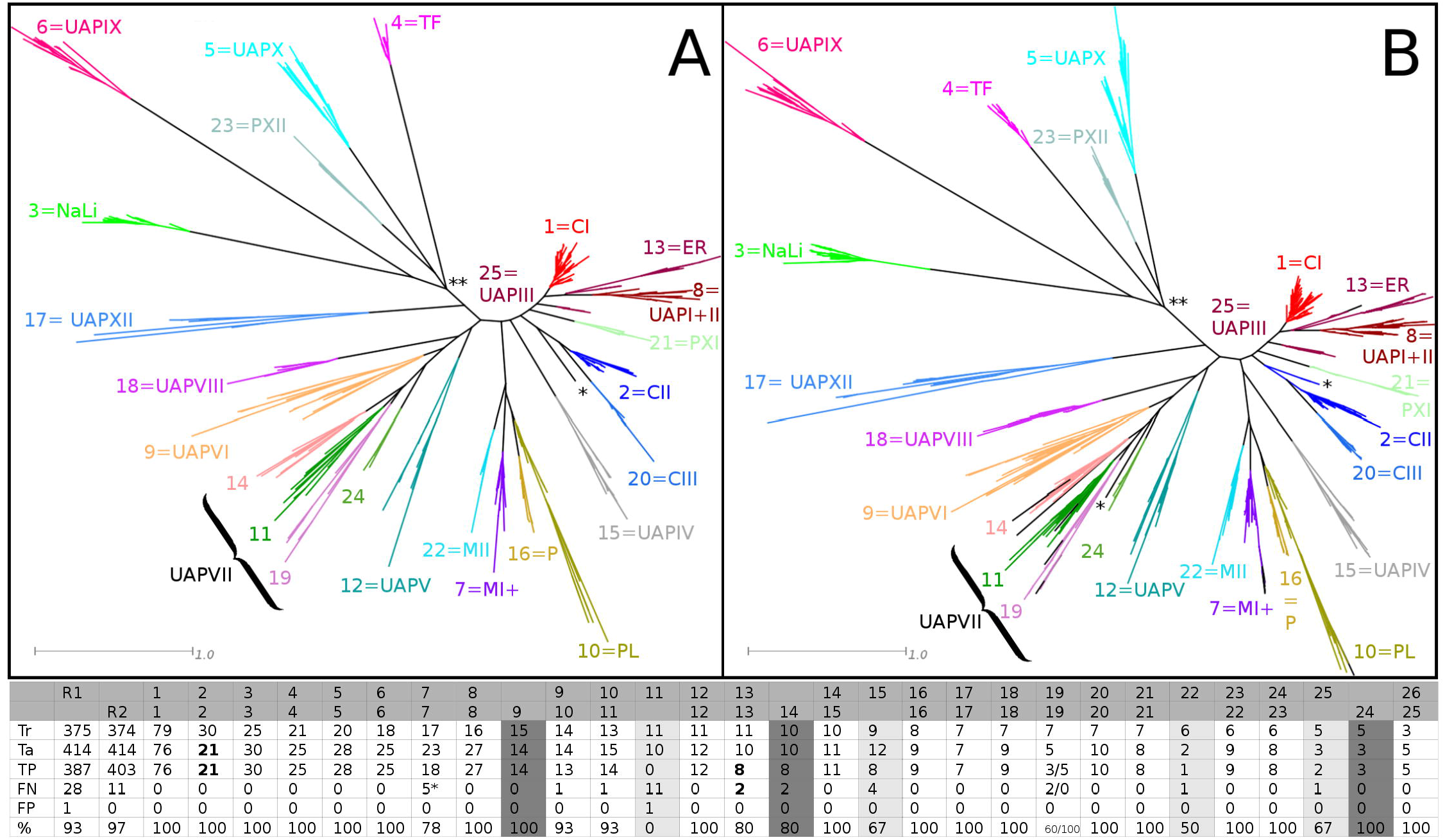
Optimized HMMERCTTER Clustering and Classification of Plant ACD Protein Superfamily. (A) Training tree with clustering; clusters numbered according to HMMERCTTER and codes applied by Bondino et al., [22]□ (MI+ combines mitochondrial I (MI) and mitochondrial-like sHSPs (ML)). (B) Final and complete reference tree with classification made using clustering shown in A. Both A and B concern the second run as detailed in the text. Colors according to HMMERCTTER output, leaves in black could not be clustered or classified. Scale bars indicate 1 amino acid substitution per site. Note that UAPVII is represented by four clusters. UAPI and II were originally identified in the final 17 proteome dataset and correspond to a single clade in the training tree. * indicates a single orphan training sequence that becomes classified in a paraphyletic clade. ** points to a local difference in tree topology that does not affect clustering and classification. The table shows the numerical results of HMMERCTTER classification. R1: 1^st^ run with specific columns in light-gray shade, R2: Optimized 2^nd^ run with specific columns in dark-gray shade. R1-G11 is R2-G14; R1-G25 is R2-G24; and R2-G9 consists of R1-G15 and R1-G22. For details see context and S Fig 2. Tr = Train; Ta = Target; TP = True Positives; FP = False Positives; FN = False Negatives; % = Coverage. Boldface numbers include paraphyletic sequences. * Concerns distant sequences as shown in S Fig. 3. Note that both the total number of train and target sequences include one orphan sequence each and that the single false positive is a training sequence.

The classification of a first run showed 93% coverage (R1 in Table of Fig 1). However, group 11 had a coverage of 0, meaning that not a single novel sequence was detected. We compared trees and analyzed *hmmsearch* output and encountered two dataset complications. First, the analysis is based on the assumption that both phylogenies are correct and as such comparable. This assumption appeared incorrect since one training sequence (VV00193000) clusters differently in both trees (S Fig 2A and B) suggesting incorrect placement in the training tree, and, as a result, incorrect HMMERCTTER clustering and poor classification. This sequence was transferred from training dataset to target dataset. Furthermore, at least one target sequence was found to contain three partial ACDs, which resulted in an elevated total score in at least two groups, which generated another classification conflict. This sequence was removed from the target dataset. We repeated the analysis resulting in an optimized clustering and classification (Fig 2). Clusters R1_C15 and R1_C22 combined with sequence VV00193000 to form a larger cluster, R2_C9, with 100% P&R in both clustering and classification. R2_C14 corresponds with cluster R1_C11 and shows 80% classification coverage. Total coverage was 97%. In general false negatives are distant sequences as exemplified by the five false negatives indicated in S Fig 3. This concerns five sequences from *Sorghum bicolor* of which four appear to derive from the same locus.

### The PG superfamily: a case showing hierarchical clustering and compositional bias

Pectin is an important structural heteropolysaccharide component of plant cell walls formed of linear chains of α-(1–4)-linked D-galacturonic acid. Rhamnose and xylose can intervene in the main chain and sugar hydroxyl groups can be substituted by methyl groups and a variety of small sugar polymers, resulting in a complex mixture of polycarbohydrates (For review see □[23]□). Plants and many of their pathogens, therefore require a number of enzymes that can degrade these polycarbohydrates, among which those of the superfamily of galacturonases (G) and polygalacturonases (PGs, SCOP identifier 51137). Many isoforms have been described and are biochemically classified according to their mode of action (exoPGs, endoPGs) and substrate specificity (PGs, rhamnoGs and xyloGs) [24]□. Then, the PG superfamily on its turn is part of the larger superfamily of pectin lyase like proteins (SCOP Identifier 51126). A number of 140 PG encoding sequences are documented in the UniProtKB/Swiss-Prot [2]□ database and used to reconstruct a Maximum Likelihood phylogeny, together forming the training dataset. The target dataset consisted of 1255 PG homolog sequences identified from EBI’s reference proteomes dataset amended with several complete proteomes from phytophagous organisms.

The training set could be clustered with 100% coverage in many ways (S Fig 4) and each clustering was used for classification of the target sequence dataset. Highest coverage (97%) was obtained using C7 (Fig 3A) (seven clusters named C7_C1 to C7-C7 and final groups C7-G1 to C7-G7), which is used as reference classification. However, the C7 clustering does not correspond perfectly with the functional clustering. C7-C2 contains two classes of exoPGs, the class of endo-xyloGs and the class of exo-rhamnoGs. C11, with C7-C2 and C7-C3 further divided into four and two subclusters respectively, does correspond with functional and hierarchical classification, albeit that exoPGs are represented by four polyphyletic clades. The C7 and C11 classifications are further discussed in detail (Fig 3).

**Fig 3:**
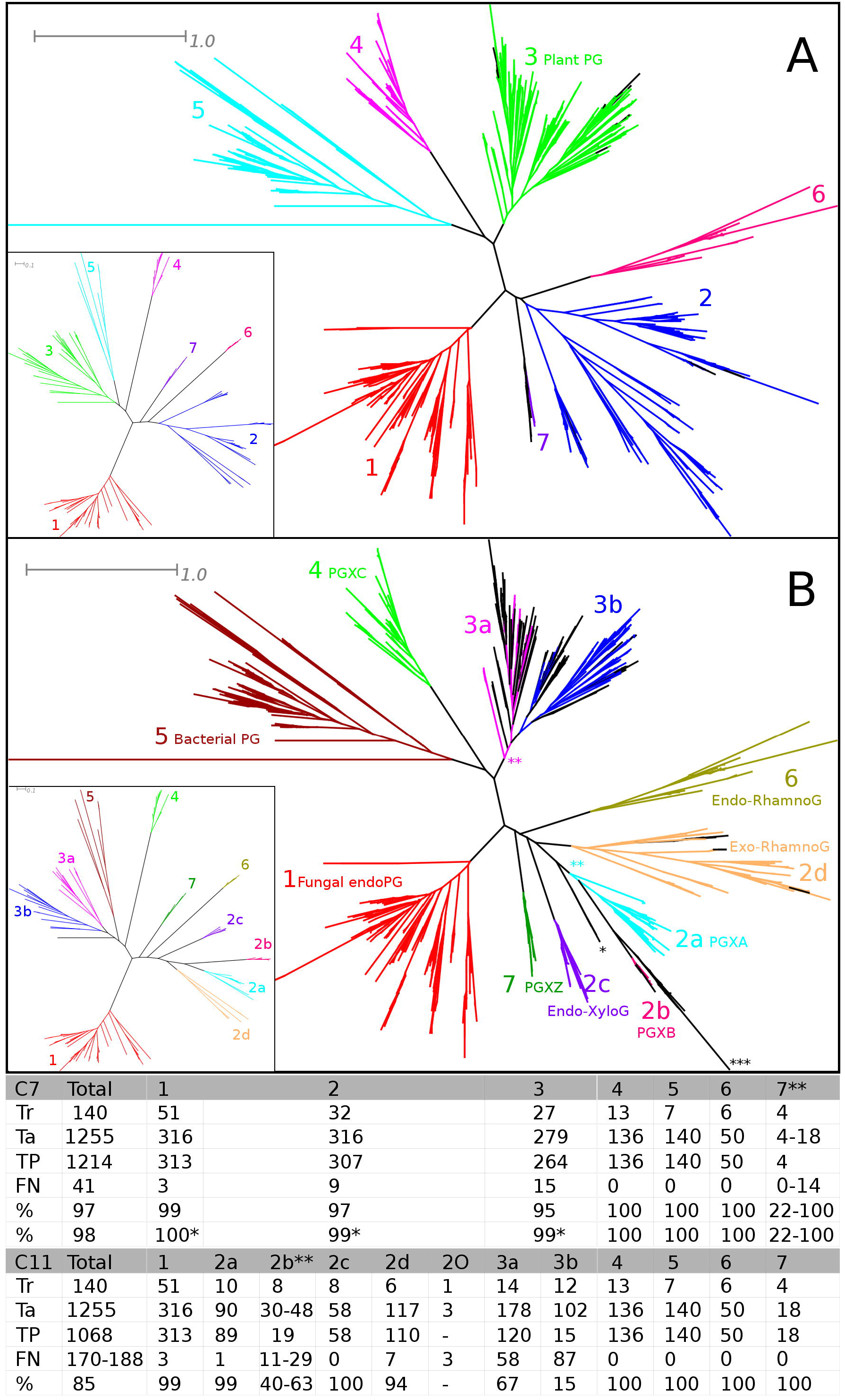
HMMERCTTER Analysis of the Polygalacturonase Superfamily. (A) C7 Clustering (inset) and classification with seven clusters. (B) C11 Clustering (inset) and classification with eleven clusters. Colors according to HMMERCTTER output, leaves in black could not be clustered or classified. Scale bars indicate 0.1 and 1 amino acid substitution per site as indicated. Functional classification is indicated in (A) for Plant PGs and (B) for the other classes where PGX stands for exo-polygalacturonase with A, B and C classes defined by Swissprot and Z corresponding to Zygomycete sequences. * Indicates a group of three orphan sequences that were not classified and could not be included in any reference classification. ** indicates instances of paraphyletic groups. *** indicates a subclade of unclassified sequences that complicate the reference classification of group 2b. Performance of the C11-2b cluster is therefore presented as an interval. The table shows the numerical results of HMMERCTTER classification. Tr = Train; Ta = Target; TP = True Positives; FN = False Negatives; % = Coverage. * Not included are 3, 7 and 11 partial sequences that appeared as false negatives in the C7 classification. ** The number of false negatives depends on the reference classification. Coverage is expressed as interval. 2O concerns the unclassifiable clade with three orphan sequences.

As expected, partitions with few clusters (e.g. C2, C3, C4 see S Fig 4A, B and C), show lower classification coverage (numbers not shown). For instance, the combined plant (C3) and bacterial (C5) PG clusters do not detect any novel sequence using the classifiers determined by partitions C2, C3 and C4. Interestingly, partitions with more clusters such as C11 (Fig 3B) and C13 (not shown), also show inferior performance.

The C7 classification performance was corrected for a small number of partial sequences from clades 1, 2 and 3 with scores slightly below the final thresholds. It also has an ambiguous reference classification. Eleven monophyletic and undetected sequences, properly detected in the C11 classification, are part of a larger monophyletic clade (S Fig 5A). Coverage interval is 22 to 100% but given the correct classification by C11, 22% should be considered as most meaningful. Strikingly, the C11-G7 score plot (S Fig 5B) shows a distinguished group with a very sharp HMMER score drop following the cut-off (from 509.3 to 269.3). The reason why other partitions yield poor classification must therefore lie in conflicting sequence identifications. Indeed, the EFRo007836 sequence, part of the clade corresponding with C11_G7 (S Fig 5A), is detected by both groups 2 and 7, and is therefore reported as conflicting sequence, arresting further analysis of either group.

### The Phospholipase C superfamily: A small, biased training set to classify a large target set

Phospholipase C (PLC) forms a class of enzymes that hydrolyze phospholipids [25]□. They are involved in cell physiology and signal transduction and there are several reasons for functional diversification, as exemplified by the fact that PLCs can have a number of different regulatory domains. Six isotypes, B, D, E, G, H and Z, are discriminated in mammals that also contain PLC-like proteins (PLC-L), which lack the second catalytic His residue [25]□. Fungi and plants also have PLCs: In tomato six isoforms have been reported [26]□ whereas in *Saccharomyces cerevisiae* only one homolog has been identified [27]□.

A total of 70 complete sequences was identified in UniProtKB/Swiss-Prot forming the training set. The target dataset consisted of 1047 sequences from EBI’s Complete Reference Proteomes dataset. The training was guided by the functional classification. Nine clusters (B, D, E, G, H, Z, L, Plant and Yeast) were assigned based on the phylogenetic clustering and UniProtKB/Swiss-Prot annotation codes that reflect the diversity. The sequence of the PLC from *Dictyostelium* (lacking any additional specification in the UniProtKB/Swiss-Prot code) was not classified and regarded as orphan sequence. Fig 4 shows the results of the classification.

**Fig 4:**
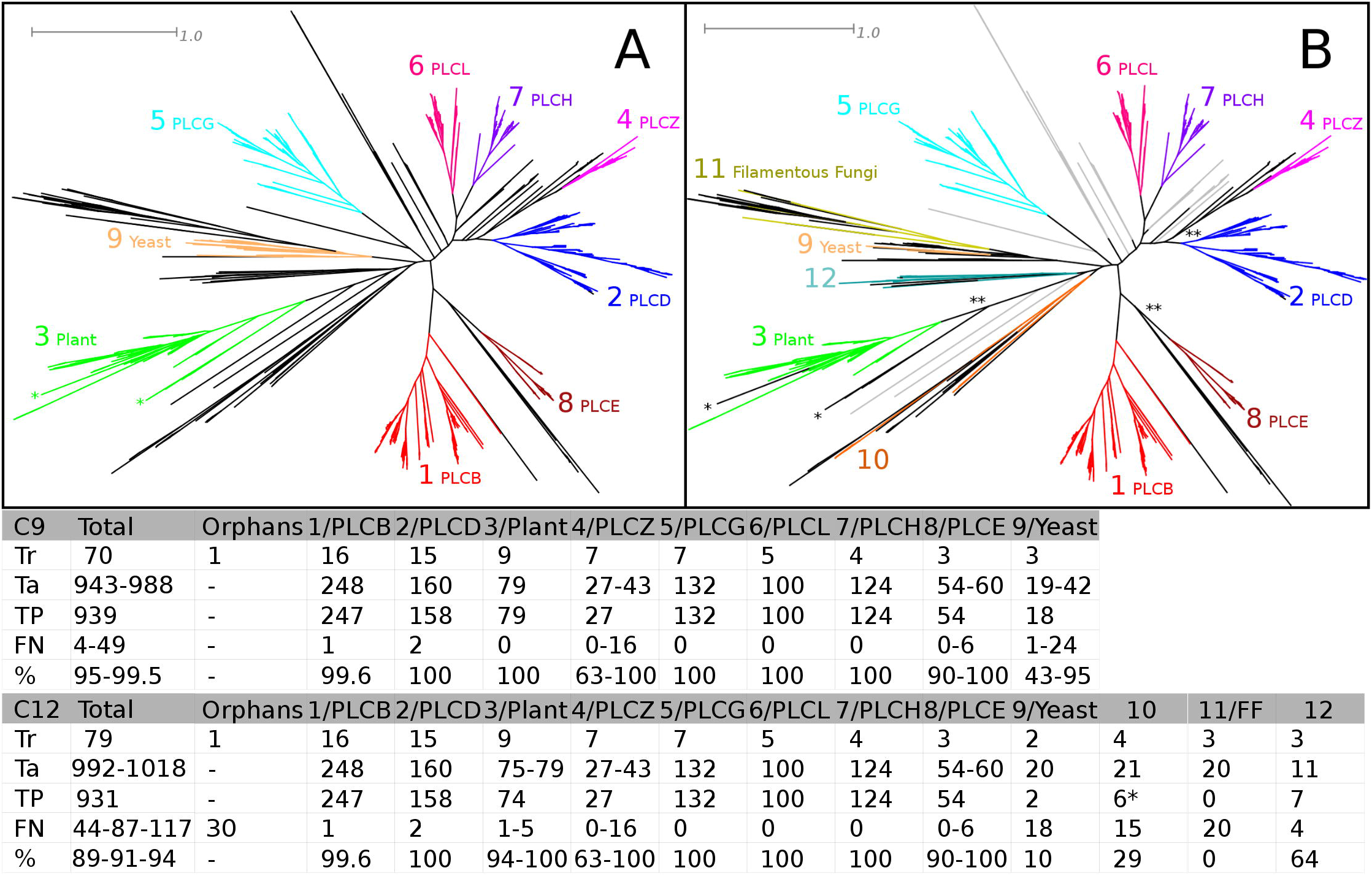
HMMERCTTER Analysis of the Phospholipase C Superfamily. (A) Initial C9 classification of nine described PLC subfamilies. (B) C12 Classification upon inclusion of additional sequences to the training set. Colors according to HMMERCTTER output of C9 clustering, leaves in black or gray could not be classified. Black leaves were included in determination of coverage or coverage interval, ** in B indicates classes where the reference classification is ambiguous. The 30 unclassifiable sequences are represented in gray. The * in A and B point to differences in the classification of Plant PLCs. The table shows the numerical results of the C9 and C12 classifications. Tr = Train; Ta = Target; TP = True Positives; FN = False Negatives; % = Coverage. * Includes the training orphan. Coverage intervals are presented when the reference classification is ambiguous and are accompanied by intervals in the number of target sequences and false negatives. The 30 unclassifiable sequences indicated in gray in B were regarded as additional false negatives resulting in the lowest coverage of the total coverage interval.

The classification of 939 sequences is nearly perfect. Strictly, only five single false negatives were identified when analyzing the classification on the reference tree (Fig 4). The performance of clusters 4 (PLCZ), 7 (Yeast) and 8 (PLCE) is ambiguous since larger clades can be considered, resulting in 49 false negatives and a coverage of 95%. However, since the dataset consisted of 1047 PLC sequences, 60 additional sequences were not classified. This is explained by the fact that the training dataset was biased: A number of clades with no training representatives is found in the final tree (See Fig 4A). We added a total of nine target sequences to the training set in order to represent three of the major unrepresented clades and repeated the analysis (Fig 4B). Surprisingly, total coverage was slightly lower at 94%. The major reason for this is the very poor performance of the yeast cluster (10%) and novel cluster 12 (0%), which contains sequences from filamentous fungi. Together they form a monophyletic clade that represents fungal PLCs, of which particularly the PLCs from filamentous fungi show high sequential divergence. Also the clade of group 10 is highly divergent and shows poor coverage (29%). The plant PLCs showed a slightly lower amount of sequences correctly identified.

## Discussion

We present and test the new protein superfamily sequence classification tool HMMERCTTER. HMMERCTTER consists of two phases: a training phase depending on a hierarchic phylogenetic clustering and a target phase in which sequences and their information are added iteratively to their classifiers, providing high sensitivity while specificity is safeguarded by imposing 100% P&R to clusters and groups as well as iteration arrest when conflicting sequence identification occurs. Here we discuss method and pipeline based on issues identified in three case studies.

### The high observed coverage is an overestimate due to an inherent lack of reference

In all three cases we found over 95% coverage of the complete datasets with groups that show 100% P&R according to the reference tree. It should be clear that the reference tree is not an ultimate benchmark dataset since *a priori* it is unknown which sequences should be considered as group member. HMMERCTTER classification performs iterative HMMER searches and includes sequences to groups while maintaining the groups at 100% P&R. Sequence classification is arrested by conflicting sequence identification or can be stopped by the user when detecting strong declines of the score drop, which follows the lowest scoring group sequence. Clearly, sequence classification terminates in the twilight zone of detection, which is inherently subject to the lack of reference. Hence, although 95% might be an overestimate, the fact that conflicting sequence classification arrests the process, at least suggests coverage is high and that HMMERCTTER is specific while it remains a high sensitivity.

### A high fidelity training set with no bias is fundamental for proper classification

The sequence incorrectly placed in the training tree of the ACD case (VV00193000, see Fig 2 and S Fig 2) had a severe impact on clustering and exemplifies the general rule that training data should be of high quality. Tree reliability is difficult to measure but in general trees with poor statistical support should be handled with care. The PLC case showed an additional training set issue. Although difficult or even impossible, bias should be avoided. HMMERCTTER is meant as a decision support system for the expert biologist, which presumably can provide a reliable training set. Still, although complete proteomes can be used as target, it is worthwhile to perform a preliminary sensitive data mining to obtain a set of target sequences restricted to possible homologs only, as we performed for the PG and PLC cases. Not only will HMMERCTTER run faster, it will also directly give an indication of performance, by which bias can be suspected, as was shown for the PLC case. Unfortunately, our attempt to correct for the bias in the PLC UniProtKB/Swiss-Prot dataset was not successful. This is at least in part due to biological complexity in the form of highly divergent subfamilies.

Orphan sequences in the training set should be avoided. They either represent incorrect sequences or result in bias since they are not included in the clustering. The ACD case had a single remaining orphan sequence. This sequence and a close homolog were classified into a group that forms a paraphyletic clade in the reference tree (Fig 2B). Paraphyletic groups were also identified in the PG classification (Fig 3B). Classification of paraphyletic sequences is possible since classification, rather than clustering, is based on HMMER profiling, basically an eloquent distance score. It should however be clear that an optimal clustering or classification corresponds with both tree topology and (known) functional classification, as was obtained for both the ACD and the PLC cases. Furthermore, the ACD case shows that sequences with repeats should be avoided. All similarity based search and classification tools inherently suffer from sequences with repeats.

### Sensitivity of individual HMMER profiles and the clustering determine P&R of the overall classification

HMMERCTTER classifies sequences using controlled iterative HMMER searches. The sensitivity of the profiles not only determines the sensitivity but also the specificity of the classification. When classification is arrested upon conflicting sequence identification, the various HMMER profiles actually compete for the unclassified sequences. In general, a HMMER profile made from a variable subfamily-MSA will be more sensitive and less specific than a HMMER profile made from a conserved subfamily-MSA. This is demonstrated by the PG case in which the partitions with few, hence, clusters with high sequence variation result in early classification conflicts of the plant and bacterial PG sequences.

On the other hand, further division of C7-C3 in C3a and C3b worsened classification, emphasizing that the final classification not only depends on the individual clusters but also on the exact clustering. This is also demonstrated by the poor classification of C7-G7, as compared to C11-G7 (See Fig 3). Here the original C7-C7 and C11-C7 clusters are identical but the additional clusters differ, resulting in different sequence identification conflict scenarios. The main difference is that in C11 the C2 cluster is subdivided into four more specific subclusters that apparently no longer detect sequences that correspond to G7. Another part of the explanation for this classification error is the fact that PGs appear to have a moderately high compositional bias, which is known to negatively affect the accuracy of HMMER scores [28]□. Similarly, convergent evolution that can be envisaged among the the four exoPG clades (2a, 2b, 6 and 7) might also negatively affect HMMER score accuracy □[28]□ and therewith specificity. The fact that the C11 clustering is capable of correctly classifying the C7 sequences, suggests that the specificities and sensitivities of the profile combinations form an important factor in determining performance.

All together this demonstrates there is a balance between group size and variability and that it is difficult to predict how well a certain clustering will perform in classification. Two aspects that will define classification performance are compactness and separateness of a cluster. Both compact (e.g. sHSP-C1 of the ACD case Fig 2) and well separated clusters (e.g. C4 from the PG case Fig 3) will show good classification. The poor classification of ACD protein clusters 11, 14, 19 and 24 (Fig 2) as well as the fungal PLCs (Fig 4) can be explained by high sequence diversity, which equals low compactness of the cluster. Sequences at larges distances will not only obtain lower *hmmsearch* scores, but will also introduce high variation into the profile made by *hmmbuild*. A divergent profile corresponds with a low specificity, which is apparently still a major limit.

On the one hand the poor classification of distant sequences shows the limit of the HMMERCTTER method, on the other hand it points to dataset issues in the form of pseudogenes or sequences derived from incorrect gene models. To the best of our knowledge no function has been assigned to any of the members of the problematic and divergent UAPVII clade. The fungal PLCs are in large part orthologs and as such not pseudogenes. Thus, biological expertise remains required. Fortunately, distant sequences will, in general, only become accepted to a group during the interactive, user-controlled part of the classification phase. In the absence of an objective method for the reliable identification of problematic sequences, the interface of HMMERCTTER’s interactive classification allows the user to use its expertise in order to make an educated decision. As such, HMMERCTTER is a decision support system.

The issue of dysfunctional sequences is problematic. Sensitive data mining, as for instance performed by the iterative JackHMMER [29]□, often results in heavily contaminated datasets, which results in severe problems while constructing an MSA. HMMERCTTER’s 100% P&R control, iteration arrest upon conflicting sequence identification and the fact that training sequences cannot be removed from the groups, prevents the inclusion of many problematic sequences and forms therefore an excellent method for sequence mining.

### Prospects

We have developed HMMERCTTER that is capable of classification of protein superfamilies sequences with both high sensitivity and specificity. This is achieved by an objective and computational approach rather than defining manually curated inclusion thresholds. The performance is high and limited mostly by aspects determined by the dataset such as training bias, errors in the training phylogeny, sequence repeats but also high sequence diversity. The 100% P&R controlled iterative approach is arrested when conflicting sequence identifications are observed. Hence, performance in the twilight zone of sequence identification is determined by a balance between sensitivity and specificity. Current efforts toward future improvements include a profound mathematical modeling of the method dedicated at properties as correctness, convergence, coverage, and measures of quality. It includes the determination of clustering quality, prediction of classification error rates, and the relationship between these two quantities.

HMMERCTTTER is a phylogenomics tool since it uses phylogeny to classify sequences. However, since it is dedicated at sub-clustering single superfamilies, it is not comparable to databases such as FunFHMM, CDD or Panther. Applying maximum likelihood phylogeny rather than a heuristic sequence based clustering will, at least theoretically, improve sub-clustering but application to large, protein space spanning databases is not feasible. However, the 100% P&R approach using HMMER appears to be applicable. The apparent error in the training phylogeny of the ACD shows that HMMER searches in a 100% P&R setting typically correspond with phylogenetic clustering, which is in correspondence with our general experience with HMMER searches. Hence, the clustering phase could be replaced by a profile database provided that the profiles and corresponding sequence database are 100% P&R. Current efforts are directed at constructing such a database that would be comparable to the aforementioned hierarchic HMMER profile databases.

## Materials and methods

### HMMERCTTER pipeline

The HMMERCTTER pipeline is written in MATLAB (The MathWorks Inc., Natick, MA, USA) and calls a number of PERL scripts that depend on Bioperl [30]□ and software packages. HMMER3 l[6]□: *hmmbuild* is used with default settings, *hmmsearch* with the option –noali. MSAs are constructed by MAFFTv7 [31]□: with the settings –anysymbol –auto. Dendroscope 3 [32]□ is used for midpoint rooting and images representing clustering on the presented phylogeny on various user interfaces.

### Datasets

The ACD training and target datasets were obtained from Bondino et al., [22]□ from which a single distant orphan sequence and the sequences from five distant subclusters were removed. PG training sequences were identified identified from UniProtKB/Swiss-Prot [2]□ by BLAST using endoPG sequence AAC64374.1 [33]□ as query. PLC training sequences were identified from Swissprot using human PLC-G sequence AAA60112.1 [34]□ as query. Target datasets for the PG and the PLC case were obtained using HMMER profiling. The PLC sequences were identified from EBI’s Reference Proteomes, which consists of 122 eukaryotic and 25 prokaryotic complete proteomes (For details see <u>*http://www.ebi.ac.uk/reference_proteomes*</u>), whereas this was amended with a number of complete proteomes from phytophagous organisms for the PG case. Sequences identified by all the superfamilies training profiles were combined and filtered by CD hit □[35]□ at 100% and scrutinized using Pfam □[7]□. In the PG case all sequences with a hit against the Glyco_hydro_28 domain were accepted, in the PLC case all sequences that hit both the PLC-X and PLC-Y domain and contain the first of two catalytic His residues were accepted. MSAs were constructed by MAFFTv7 [31]□ using the slow iterative global refinement (FFT-NS-i) mode for PGs and the multiple domain iteration (E-INS-i) mode for PLCs and subsequently corrected by Rascal [26]□. PHYML3 [36]□ using the LG model was used for phylogenetic tree reconstruction following BMGE [37]□ trimming with BLOSUM62 matrix and an entropy cut-off of 0.9. Complete trees were constructed with all identified sequences. The training tree of the second PLC analysis is an excerpt of the PLC reference tree. A package containing source code and full dataset will be deposited at Github and is available for reviewers at: https://www.dropbox.com/s/aacao6ggcak30bg/Repo.tar.gz?dl=0

## Legends Supplemental

**Files S Fig 1: HMMERCTTER User Interfaces**

Shown are single examples obtained from the training (A) and the target phase (B). Blue asterisks are scores of sequences of the complete dataset (either training or combined), green circles are scores of sequences of the cluster or group in question. The red line shows the drop in scores among successive sequences, also indicated as score drop. The magenta line shows the current threshold and can be moved in order to determine the threshold in the interactive part of the classification. In the training the user must accept or reject the 100% P&R cluster. During classification the user can accept the cluster, return to the former or initial state or add seemingly negatives as indicated for posterior 100% P&R testing.

**S Fig 2: Incongruent Clustering of ACD Train Tree severely affects Classification**

(A) Detail of poor classification of first run using clustering based on train-tree with sequence VV00193000 clustered differently than in the shown final tree. Sequences of GR1-11 including VV00193000 in red; GR1-15 in gray nested inside the clade containing GR1-22 light green lines. Target sequence ME22590va contains three ACDs also generating classification conflicts. (B) Detail of improved classification of second run obtained upon removal of VV00193000 and ME22590va from training and target set, respectively. GR2-14 shows 80% coverage, formerly 0% with G1-11. GR2-9 shows 100% coverage, which is an improvement of the 67 and 50 of constituents GR1-15 and GR1-22. Numbers also in Fig 1.

**S Fig 3: Sequences at large Distances impede Classification Coverage.**

Detail of cluster 7/M+ demonstrating that particularly sequences at large distances (ML) are often not detected. The indicated sequences without shading are training sequences whereas the sequences in red shade are false negatives. The arrow points to a monocotyledon sub clade.

**S Fig 4: C2, C3 and C4 Clustering and Classification of the Polygalacturonase Superfamily**

(A1) C2 Clustering; (A2) C2 Classification; (B1) C3 Clustering; (B2) C3 Classification; (C1) C4 Clustering; (C2) C4 Classification. Colors according to HMMERCTTER output, leaves in black could not be clustered or classified. Scale bars indicate 1 amino acid substitution per site.

**S Fig 5: Poor Classification of Group 7 by C7 Clustering**

(A) Classification of group 7 by C7 clustering. In purple sequences from C7-G7 group. Other leaves represent sequences identified by the C11 clustering. EFro007836 is identified by both C7-C2 and C7-C7, preventing classification. (B) Score plot of C11-G7. Blue asterisks are scores of sequences of the complete dataset, green circles are scores of sequences group C11-G7. The red line shows the drop in scores among successive sequences. The magenta line shows the current threshold.

